# Meta-analysis of immune induced gene expression changes in diverse *Drosophila melanogaster* innate immune responses

**DOI:** 10.1101/2021.09.23.461556

**Authors:** Ashley L. Waring, Joshua Hill, Brooke M. Allen, Nicholas M. Bretz, Nguyen Le, Pooja Kr, Dakota Fuss, Nathan T. Mortimer

**Author notes:** Correspondence: Nathan T. Mortimer, School of Biological Sciences, Campus Box 4120, Illinois State University, Normal, IL 61790, USA.

## Abstract

**Background:** Organisms are commonly infected by a diverse array of pathogen types including bacteria, fungi, viruses, and parasites, and mount functionally distinct responses to each of these varied immune challenges. Host immune responses are characterized by the induction of gene expression in response to infection. However, the extent to which expression changes are shared among responses to distinct pathogens is largely unknown.

**Results:** We performed meta-analysis of gene expression data collected from *Drosophila melanogaster* following infection with a wide array of pathogens. We identified 62 genes that are significantly induced by infection. While many of these infection-induced genes encode known immune response factors, we also identified 21 genes that have not been previously associated with host immunity. Examination of the upstream flanking sequences of the infection-induced genes lead to the identification of two conserved enhancer sites. These sites correspond to conserved binding sites for GATA and nuclear factor κB (NFκB) family transcription factors and are associated with higher levels of transcript induction. We further identified 31 genes with predicted functions in metabolism and organismal development that are significantly downregulated following infection by diverse pathogens.

**Conclusions:** Our study identifies conserved gene expression changes in *Drosophila melanogaster* following infection with varied pathogens, and transcription factor families that may regulate this immune induction. These findings provide new insight into transcriptional changes that accompany *Drosophila* immunity. They may suggest possible roles for the differentially regulated genes in innate immune responses to diverse classes of pathogens, and serve to identify candidate genes for further empirical study of these processes.

## Background

Organisms encounter a broad range of pathogens in their natural environments and have evolved immune defenses that allow them to survive infection by diverse pathogen classes. Most metazoan hosts make use of a highly conserved suite of innate immune responses to defend against pathogen infection [1]. These responses are typified by a multi-step process, beginning with pathogen recognition and resulting in activation of the appropriate immune mechanism [2]. This immune activation is accompanied by changes in gene expression, including both the induction of immune response transcripts and the downregulation of transcripts encoding proteins with non-immune functions [3, 4]. The induced immune gene products then play a role in either eliminating the pathogen (pathogen resistance) or allowing the organism to survive despite infection (pathogen tolerance). Differential gene expression analysis has proven to be a valuable approach to uncover the genetic basis for a variety of traits [5–8], including the immune response to infection [9–11]. These studies have revealed the importance of conserved immune signaling pathways including Toll, Immune deficiency (IMD) and JAK-STAT across diverse organisms [12–19].

While hosts may encounter numerous distinct pathogenic species, the pathogens are often categorized into four general classes: bacterial pathogens, fungal pathogens, viruses, and parasites [20, 21]. Innate immune responses can be highly specific to the category of pathogen encountered and may include both cellular and humoral mechanisms [2, 22]. For instance, bacterial infection is often countered by the production of secreted antimicrobial peptides (AMPs) and immune cell mediated phagocytosis of the invading microbes [23, 24]. Alternatively, antiviral immunity may include distinct features such as RNA interference or the recognition and cytolysis of infected host cells [25, 26]. Highly conserved transcriptional signatures in response to distinct pathogen types have been uncovered, but less is known about common patterns of gene induction against multiple pathogens [27–30].

The genetic model organism *Drosophila melanogaster* is a commonly used and powerful system to understand conserved innate immune processes [31–33]. Like other animals, *Drosophila* mount specific responses to infection by distinct pathogen classes [20, 22]. Numerous studies have investigated changes in gene expression in *Drosophila* hosts following infection by a wide range of pathogens including multiple species of bacterial and fungal pathogens, viruses, and parasitoid wasps (Table 1) [18, 19, 34–40]. These studies provide a unique opportunity for the comparative analysis of immune responses by a single host organism against diverse pathogens. A useful method to perform comparative analyses is the meta-analysis of gene expression studies. Such meta-analyses attempt to directly compare results from multiple studies while controlling for inter-study differences [41]. Here, we use a common meta-analysis approach to perform a comparative analysis on multiple previously described *Drosophila* infection studies (Table 1) to identify genes whose expression are similarly altered across infection by distinct pathogen classes.

**Table 1.** List of datasets used in the meta-analysis.

| GEO Accession | Pathogen Type | Pathogen | Host Stage | Reference |
| --- | --- | --- | --- | --- |
| - | Bacteria | *Escherichia coli* + *Micrococcus luteus* | Adult | [34] |
| - | Bacteria | *E. coli*, *M. luteus* + *Enterococcus faecalis* | Adult | [18] |
| GSE37708 | Bacteria | *E. coli* | Adult | [35] |
| GSE5489 | Bacteria | *E. coli* | Larva | [36] |
| - | Fungus | *Beauveria bassiana* | Adult | [34] |
| - | Fungus | *Aspergillus fumigatus* | Adult | [18] |
| GSE2828 | Virus | *Drosophila C Virus* | Adult | [37] |
| GSE42726 | Virus | Sindbis virus (transgenic *Drosophila* model) | Adult | [38] |
| GSE31542 | Virus | Flock House Virus | Adult | [39] |
| GSE31542 | Virus | Sindbis virus | Adult | [39] |
| GSE25522 | Parasite | *Ganaspis xanthopoda* | Larva | [40] |
| GSE8938 | Parasite | *Leptopilina boulardi* | Larva | [19] |

## Results

### Meta-analysis of genome-wide transcript levels following pathogen infection reveals a signature of differential gene expression

We performed a meta-analysis on 10,818 genes across 12 gene expression studies following infection by a variety of pathogens (listed in Table 1). To identify genes showing significant expression changes across these studies, we used the non-parametric rank products approach with an estimated percentage of false prediction (pfp) threshold of 0.05. In this method, the observed fold-change of each gene is ranked within each study and the rank-product of each gene is calculated as the geometric mean of the ranks of a given gene across all of the studies. Genes with rank-products that significantly differ from a uniform distribution are considered to be up- or down-regulated [42, 43]. The use of ranks, rather than experimental values, makes this approach robust to differences between experimental platforms and allows for the comparison between multiple studies [43, 44]. Using this approach, we identified 62 genes that were induced across these infection conditions (Additional File 1) with an average log_2_ fold change (logFC) of 1.17, and a logFC range of 0.62 to 2.27. We further identified 31 genes that were significantly downregulated across these infection conditions (Additional File 2). These downregulated genes have an average logFC of - 0.75, with a logFC range of −0.26 to −1.40.

The identified genes were mapped onto their chromosomal locations (Figure 1) and found to be distributed throughout the autosomal chromosome arms, with few genes mapping to the X chromosome and none on chromosome 4. Given that only ~80 genes of ~18,000 total genes in the *D. melanogaster* genome are found chromosome 4 [45, 46], the lack of immune regulated genes on chromosome 4 is unsurprising. On the other hand, the apparent lack of immune regulated genes on the X chromosome was unanticipated. We therefore used 2-sample proportion tests to assay the distribution of genes on each chromosome arm (Table 2). We find that induced genes are significantly enriched (X^2^ = 8.74, p = 0.0031), and that downregulated genes are significantly under-represented (X^2^= 4.56, p = 0.033), on chromosome 2R. Additionally both induced and downregulated genes are under-represented on the X chromosome (induced: X^2^= 3.88, p = 0.049; downregulated: χ^2^= 11.13, p = 8.5×10^−4^). This relative lack of genes was unexpected given the presence of numerous immune response genes on the X chromosome, although interestingly, the antimicrobial peptide class of immune effectors is also under-represented on the X chromosome [47].

**Figure 1.**
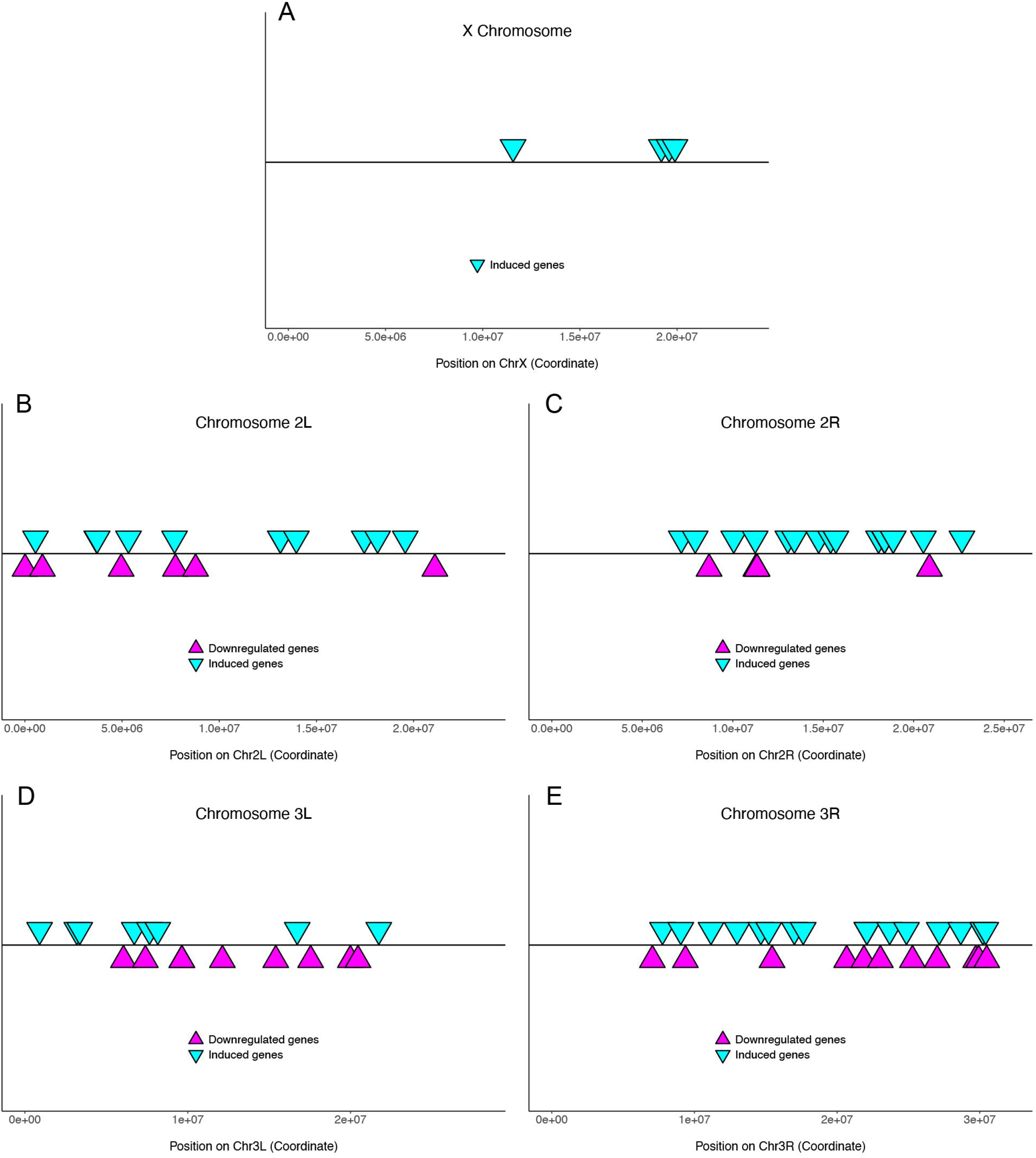
Chromosomal location of altered genes. Each identified gene has been mapped to its chromosomal location, indicated by its position on each chromosome arm of the *Drosophila melanogaster* genome (A-E). For each panel, the x axis represents the genomic position, inverted cyan triangles indicate the positions of induced genes and the magenta triangles indicate the positions of downregulated genes.

**Table 2.** Distribution of analyzed genes across chromosome arms. The Dataset category contains the genes that were measured in all 12 datasets. ^a^ enriched relative to Dataset control, ^b^ under-represented relative to Dataset control; determined by p < 0.05 from 2-sample test for equality of proportions with continuity correction.

| Sample | X | 2L | 2R | 3L | 3R | 4 | U | Total |
| --- | --- | --- | --- | --- | --- | --- | --- | --- |
| Dataset | 1793 | 1951 | 2129 | 2103 | 2734 | 65 | 43 | 10818 |
| Induced genes | 4<sup>b</sup> | 11 | 22<sup>a</sup> | 8 | 17 | 0 | 0 | 62 |
| Downregulated genes | 0<sup>b</sup> | 6 | 5<sup>b</sup> | 9 | 11 | 0 | 0 | 31 |

Next, we assayed the distribution of the identified genes within each chromosome arm for overall uniformity and the presence of gene clusters. Using the Kolmogorov-Smirnov test of uniformity, we find that the genes are evenly distributed along chromosomes (Table 3), but this analysis did identify the presence of 7 gene clusters. This represents a significant degree of clustering compared to background controls (induced p = 1.76 x 10^−7^; downregulated: p = 0.003). We identified 5 clusters of induced genes (annotated in Additional File 1) including the Bomanin family gene clusters (found on chromosomes 2R and 3R), clusters comprising the Diptericin (chromosome 2R) and Cecropin (chromosome 3R) antimicrobial peptide families, and a cluster of two unstudied genes on chromosome 2L (*CG9928* and *CG16978*). We also identified 2 clusters of downregulated genes (annotated in Additional File 2) including a cluster of Trypsin genes (chromosome 2R) and a cluster of predicted S1A family serine protease genes (*CG18179* and *CG18180*; chromosome 3L).

**Table 3.** Uniformity of altered genes within chromosome arms. The uniformities of induced and downregulated genes within each chromosome arm were independently assessed by Kolmogorov-Smirnov test. *D* is the calculated Kolmogorov-Smirnov distance, Up = induced (upregulated) genes, Down = downregulated genes.

| Chromosome Arm | *D* Up | P value Up | *D* Down | P value Down |
| --- | --- | --- | --- | --- |
| X | 0.62 | 0.05 | - | - |
| 2L | 0.17 | 0.87 | 0.41 | 0.20 |
| 2R | 0.24 | 0.15 | 0.51 | 0.10 |
| 3L | 0.41 | 0.10 | 0.25 | 0.52 |
| 3R | 0.17 | 0.66 | 0.31 | 0.19 |

### Numerous infection-induced genes play identified roles in host immunity

Our meta-analysis identified 62 genes that are significantly upregulated following infection. Of these, 42 have been previously linked with host immunity (Table 4). This list includes genes that have been previously implicated in resistance to each of the pathogen categories. The list also includes genes with membership in the Toll, IMD, JAK-STAT and Jun N-terminal kinase (JNK) conserved immune signaling pathways. Accordingly, Gene Ontology (GO) term analysis revealed that the immune induced genes are enriched in multiple biological processes linked to responses to external stimuli including immune response (GO:0009655), response to biotic stimulus (GO:0009607), response to wounding (GO:0009611), and response to stress (GO:0006950) (Figure 2, Additional File 3).

**Table 4.** Infection induced genes with previous links to immune function or immune signaling pathways.

| Gene Name | Function | Immune Pathway | References |
| --- | --- | --- | --- |
| *AttA* | Antimicrobial peptide | IMD | [116, 117] |
| *AttD* | Antimicrobial peptide | IMD | [116] |
| *Bbd* | Production of AMP-like peptides | Toll | [51] |
| *BomBc1* | AMP-like | Toll | [49, 50] |
| *BomBc2* | AMP-like | Toll | [49, 50] |
| *BomBc3* | AMP-like | Toll | [49, 50] |
| *BomS1* | AMP-like | Toll | [49, 50, 118] |
| *BomS2* | AMP-like | Toll | [49, 50, 118] |
| *BomS3* | AMP-like | Toll | [49, 50, 118] |
| *BomS5* | AMP-like | Toll | [49, 50, 118] |
| *BomS6* | AMP-like | Toll | [49, 50, 118] |
| *BomT2* | AMP-like | Toll | [49, 50] |
| *BomT3* | AMP-like | Toll | [49, 50] |
| *CecB* | Antimicrobial peptide | IMD | [119, 120] |
| *CecC* | Antimicrobial peptide | IMD | [119, 121] |
| *Def* | Antimicrobial peptide | IMD, Toll | [18, 122] |
| *DptA* | Antimicrobial peptide | IMD | [119, 123] |
| *DptB* | Antimicrobial peptide | IMD | [119, 124] |
| *Drs* | Antimicrobial peptide | Toll | [125, 126] |
| *Ets21C* | Transcription factor | IMD | [127] |
| *BaraA* | Antimicrobial peptide | Toll | [128] |
| *Irc* | Oxidant detoxification | - | [129] |
| *lectin-24A* | Carbohydrate binding | - | [130] |
| *Listericin* | Antimicrobial peptide | JAK-STAT | [131] |
| *mat* | - | JAK-STAT | [132, 133] |
| *Mtk* | Antimicrobial peptide | IMD, Toll | [134, 135] |
| *nec* | Serpin | Toll | [136] |
| *NimB1* | Pathogen recognition (predicted) | - | [137, 138] |
| *PGRP-SA* | Pathogen recognition | Toll | [53, 139] |
| *PGRP-SB1* | Antimicrobial effector | IMD | [140, 141] |
| *PGRP-SD* | Pathogen recognition | Toll | [142] |
| *Rel* | Transcription factor | IMD | [70, 143] |
| *Sid* | DNA endonuclease | Toll | [18, 144] |
| *Sp7* | S1A Serine Protease | Toll | [145, 146] |
| *SPE* | S1A Serine Protease | Toll | [54, 147] |
| *Tep2* | Thioester-containing Protein | Toll | [148] |
| *TotM* | - | JAK-STAT | [149, 150] |
| *CG13675* | Chitin Binding | IMD | [151] |
| *CG14957* | Chitin Binding | JNK | [133] |
| *CG3505* | S1A Serine Protease | Toll/IMD | [152] |
| *CG5909* | S1A Serine Protease | Toll/IMD | [18] |

**Figure 2.**
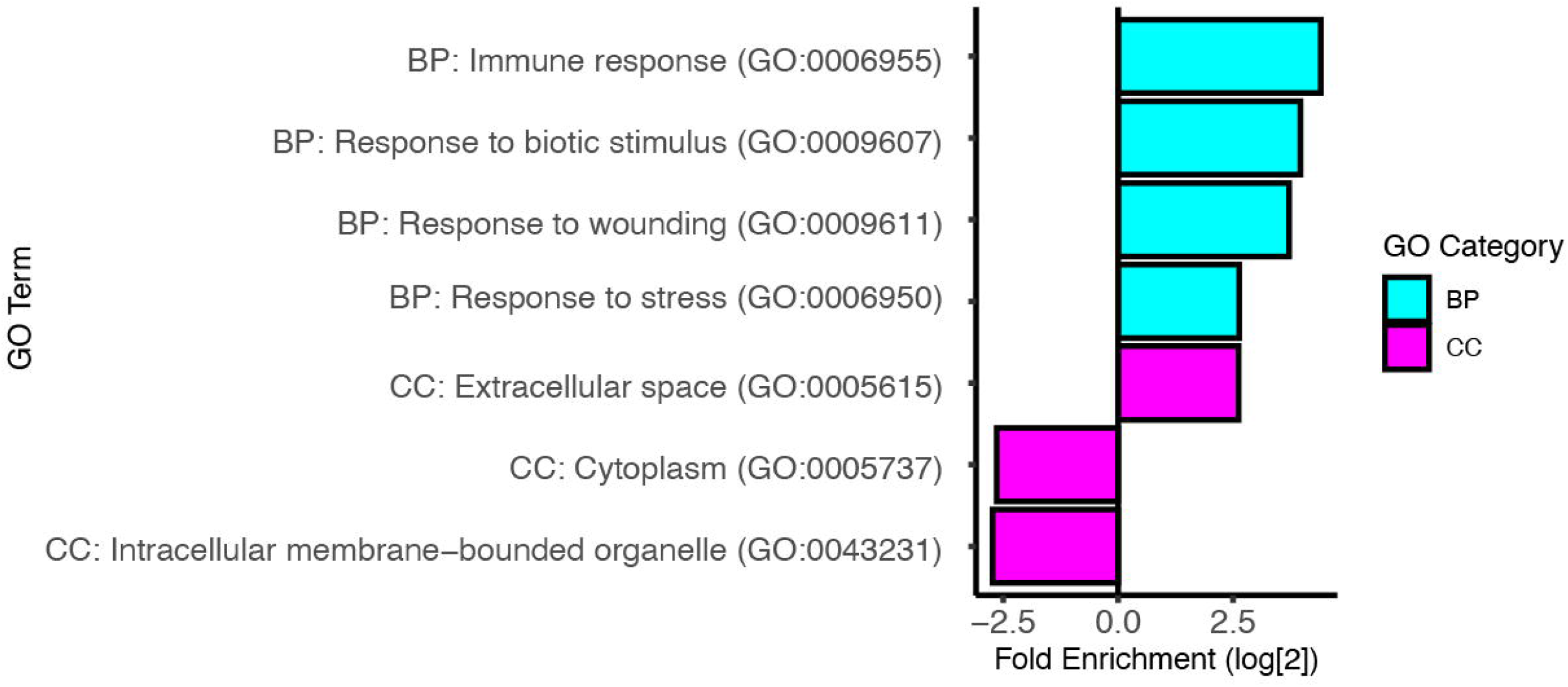
Gene ontology analysis of infection induced genes. The log_2_ fold enrichment for selected GO terms. The Biological Process (BP) category is shown in cyan, and the Cellular Component (CC) category is shown in magenta. See Additional File 3 for complete GO term analysis of induced genes.

Our meta-analysis results suggest that infection by a broad range of pathogens can lead to differential regulation of immune signaling pathways. We found that several genes implicated in pathogen recognition (*NimB1*, *lectin-24A* and *Tep2*), along with regulators of the Toll (*nec, PGRP-SA, SPE* and *Sp7*) and IMD (*PGRP-SD* and *Rel*) pathways, are induced following infection. We also identified a broad range of immune effector molecules including antimicrobial peptides (AMPs) and AMP-like immune induced genes. The *D. melanogaster* genome encodes numerous AMP families [48], and we identify members of nearly all of these families including the Attacins (AttA and AttD), Cecropins (CecB and CecC), Defensin, Diptericins (DptA and DptB), Drosomycin, IMPPP/Baramycin A, Listericin, and Metchnikowin (Table 4). This broad induction is particularly interesting given that the Toll and IMD signaling pathways and many of these AMP/AMP-like families have distinct pathogen targets [24].

Members of the *Bomanin* (*Bom*) AMP-like gene family have been shown to act downstream of Toll pathway signaling in antimicrobial immunity [49, 50]. We identified 10 of the 12 *Bom* family genes as induced in our meta-analysis (Table 4); of the other 2 *Bom* genes, *BomS4* is significantly induced following parasite infection, but not following infection by the other pathogens, and *BomT1* is not represented in our dataset. *Bom* genes are found in the genome in two clusters, and we identified genes from both clusters in our analysis. We additionally identified the *bombardier* (*bbd*) gene as induced in our analysis (Table 4). Like the *Bom* genes, *bbd* also acts in antimicrobial immunity downstream of Toll, and *bbd* mutants fail to produce the short-form class of *Bom* peptides (*BomS*) [51]. The finding that *bbd* is induced alongside *Bom* genes lends further support for a role for *Bom* family activity following infection.

### Infection induced genes encode proteins with a variety of predicted functions

The proteins encoded by the infection induced genes have a wide array of predicted molecular functions. This includes both immune associated functions like antimicrobial activity and peptidoglycan recognition, and a variety of other functions such as ion transport (*MFS12*), deoxyribonuclease activity (*Sid*), and acyl transferase activity (*CG14219*). Notably, we identified 5 members of the S1A protease family (*SPE*, *Sp7*, *CG3505*, *CG18563* and *CG5909*). The S1A family is comprised of more than 200 genes and includes both active proteases and catalytically inactive protease homologs [52]. S1A family members have been previously linked to immune responses against a variety of pathogens [53–57]. Due to the wide array of encoded protein activities, our GO term analysis did not identify any significant enrichment for molecular function. However, we did identify an enrichment of genes encoding proteins that are secreted into the extracellular space (GO:0005615) and an under-representation of genes encoding cytosolic (GO:0005737) and intracellular membrane-bounded organelle localized proteins (GO:0043231) (Figure 2, Additional File 3).

### Motif finding suggests that the infection induced genes are regulated by NFκB and GATA family transcription factors

The timing and levels of mRNA transcription are tightly regulated by the activity of a wide array of transcription factors. Multiple transcription factor families, including the nuclear factor κB (NFκB), nuclear factor of activated T cells (NFAT), signal transducer and activator of transcription (STAT) and erythroblast transformation specific (ETS) factors have been linked to transcription induction following infection [58–61]. We predicted that the induced genes uncovered in our meta-analysis are co-transcriptionally regulated, and share a common set of transcription factors. To test this prediction, we analyzed the 250 bp of genomic sequence upstream of the annotated transcription start site of each of the induced genes using the iMotifs *de novo* motif finder [62]. We reasoned that these sequences likely included the promoter and proximal enhancers for each gene, and our approach would allow us to uncover any motif that was found in the upstream region of the majority of the induced genes.

Our analysis led to the identification of 3 consensus motifs (Figure 3A-C). These consensus sequences were searched against the complete database of known *D. melanogaster* transcription factor binding sites using the Tomtom web server [63]. We found that our Motif 1 showed significant similarity to the binding site for the GATA factor *GATAe* (p = 6.0 x 10^−6^; Figure 3A), and that our Motif 2 showed significant similarity to the binding site for the NFκB factor *dl* (p = 5.6 x 10^−4^; Figure 3B). It can be challenging to correctly identify the binding site for a specific member for either of these transcription factor families [64, 65], and so we will refer to the sites by their general classifications as GATA and NFκB sites for Motifs 1 and 2, respectively. Motif 3 did not show significant similarity to known *D. melanogaster* transcription factor binding sites. However, its core sequence matches the TATA box characteristic of many eukaryotic core promoters (Figure 3C) [66, 67]. This finding supports our use of upstream genomic sequences to capture gene promoter regions.

**Figure 3.**
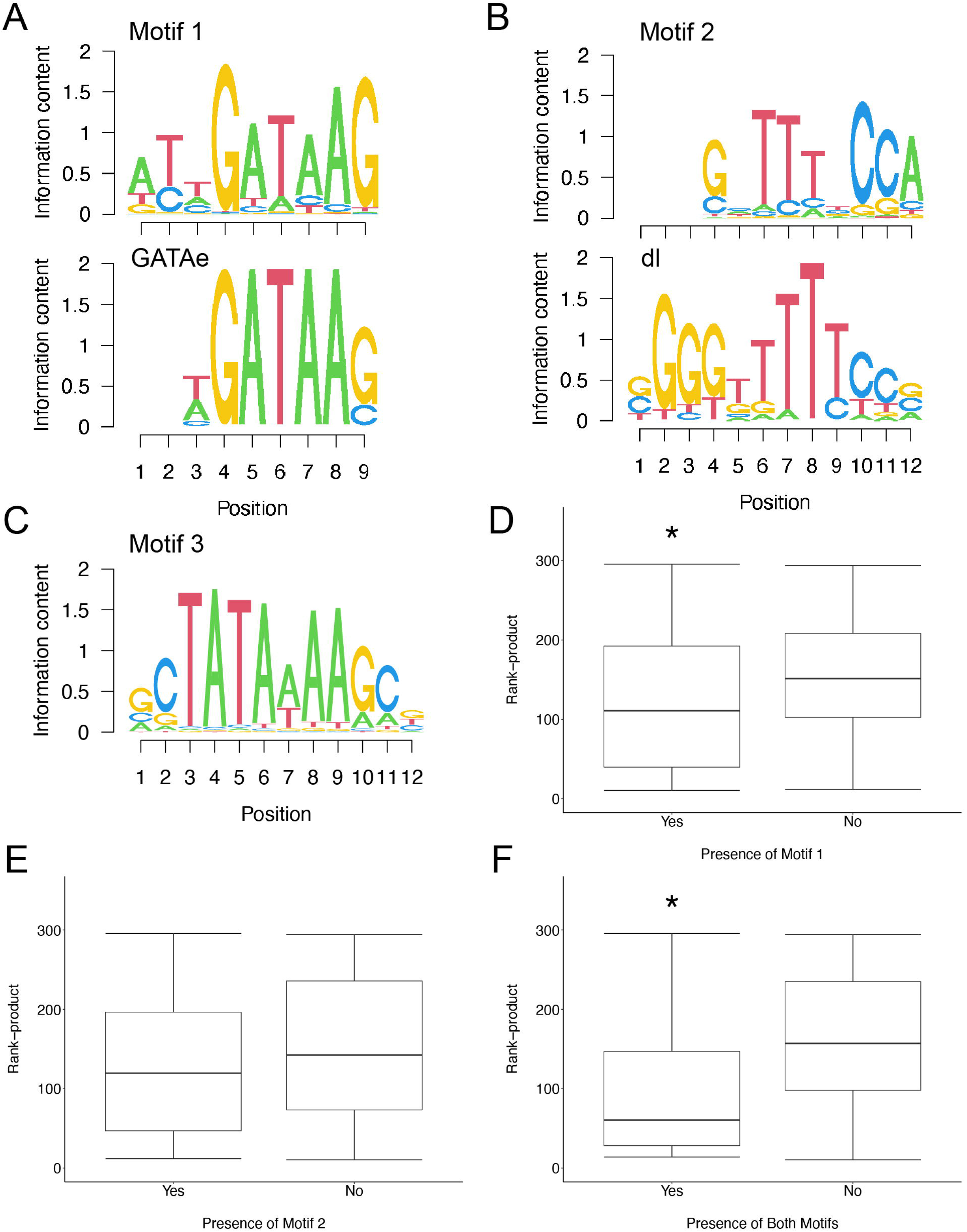
Motifs associated with infection induced genes. *De novo* motif finding identified 3 motifs (Motifs 1-3) that are enriched in the upstream sequences of the induced genes. The consensus motifs are represented as sequence logos (A-C, top). Motif matching identifies Motif 1 as being significantly similar to the GATAe binding site (A). Motif 2 shows significant similarity to the dl binding site (B). (D-F) Box-whisker plots showing the distribution of rank-products for induced genes with and without the indicated motifs. A lower rank-product is indicative of higher expression levels. (D) Induced genes with Motif 1 in the upstream region have significantly lower rank-products. (E) The presence of Motif 2 does not impact the rank-product distribution. (F) Induced genes with both motifs have significantly lower rank-products. Asterisk (*) indicates p < 0.05 relative to induced genes without the indicated motif.

Since both GATA and NFκB factors have been previously linked to infection induced transcription in *Drosophila* [68–71], we next tested whether these sites are more common in the upstream regions of our induced genes than of a control list of unchanged genes from the meta-analysis (Additional File 4). We find that our identified induced genes are significantly more likely that the background control to contain either a Motif 1/GATA or Motif 2/NFκB site (Fisher’s Exact Test odds ratio 7.5, p = 1.6 x10^−6^). More specifically, both Motif 1 and Motif 2 are enriched in the upstream regions of our induced genes (Motif 1: odds ratio 4.7, p = 1.0 x 10^−4^; Motif 2: odds ratio 5.2, p = 2.5 x10^−5^; Table 5), and the induced genes are also significantly more likely to contain both motifs (odds ratio 16.2, p = 5.3 x10^−6^; Table 5).

**Table 5.** Enrichment of predicted transcription factor binding sites in the induced genes compared to unchanged control genes. * p > 0.05 compared to unchanged control by Fisher’s exact test.

| Gene Set | Motif 1 | Motif 2 | Both | Neither | Total |
| --- | --- | --- | --- | --- | --- |
| Induced | 36* | 39* | 22* | 9* | 62 |
| Unchanged | 14 | 15 | 2 | 35 | 62 |

While this enrichment is suggestive that GATA and NFκB factors have an impact on immune induced expression, we wanted to test this hypothesis more explicitly. We used the Wilcoxon rank sum exact test to test the effect of each site on the rank-product of the induced genes. We would predict that if the presence of Motif 1/GATA or Motif 2/NFκB (or both) sites has a positive effect on expression, then transcripts with these sites would have significantly lower rank-products (indicative of higher expression). We find that genes with Motif 1 have significantly lower rank-products that those without (W = 595, p = 0.035; Figure 3D), while the presence of Motif 2 alone has little to no impact on rank-product (W = 510, p = 0.189; Figure 3E). In contrast, transcripts with both motifs have significantly lower rank-products (W = 616, p = 4.53 x10^−3^; Figure 3F), with the strength of this effect hinting at a possible synergistic effect of GATA and NFκB factors.

### Transcripts that are downregulated following infection may contribute to metabolic changes and life history tradeoffs observed with immune activation

Our meta-analysis identified 31 genes that are significantly downregulated following infection with the various pathogens (Additional File 2). Most of these genes may be predicted to be influenced by life history tradeoffs that occur following infection. Mounting an immune response is energetically costly, and following infection organismal metabolism is altered [72–74]. We find a wide range of genes linked to metabolism are downregulated following infection including genes linked to amino acid metabolism (*Lsp1*β*, Lsp2, Srr*), lipid metabolism (*CG17192, mag*), and carbohydrate metabolism (*LManVI, Sodh-1*). The shifting of resources towards immunity is often at the expense of organismal development or fertility [75–77], and among our downregulated genes, we find genes associated with these processes including *fln, Act88F, CG33259, Cpr92F,* and *TpnC47D*.

In many organisms, pathogen infection leads to coordinated changes in host physiology and behaviour, known as sickness behaviour [78]. These changes include decreased host appetite and feeding following infection in a wide range of host species, including *Drosophila* [78–82]. Accordingly, we find that genes involved in feeding behaviour (*fit*) and nutritional stress (*CG18179, CG18180*), along with putative digestive enzymes in the Trypsin and Jonah protease families are all downregulated following pathogen infection [83–85]. Interestingly, despite the widespread prevalence of infection induced anorexia, this mechanism is not uniformly protective, and instead can lower host resistance to certain pathogens [82, 86, 87].

Our GO term analysis (Figure 4, Additional File 4) of these downregulated genes reveals an enrichment in genes involved in proteolysis (GO:0006508), and specifically in serine-type peptidase activity (GO:0008236). The enrichment in genes found in the larval serum protein complex (GO:0005616) likely reflects the observed metabolic change. Like with the induced genes, we also find an under-representation of genes encoding proteins that localize to intracellular membrane-bounded organelles (GO:0043231).

**Figure 4.**
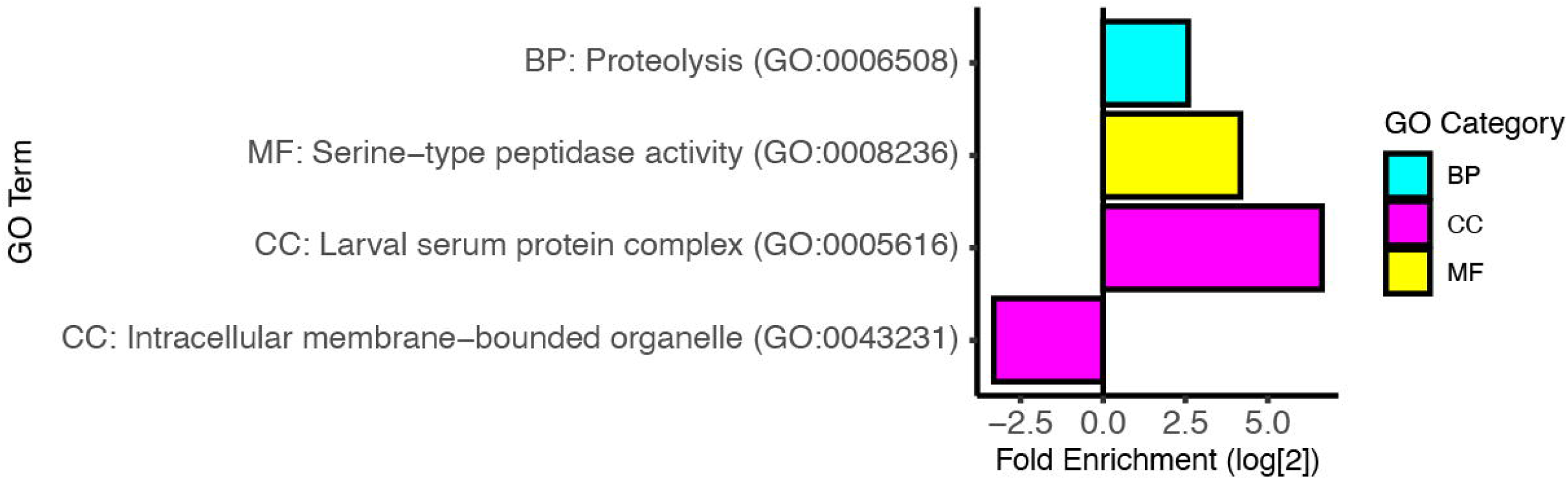
Gene ontology analysis of downregulated genes. The log_2_ fold enrichment for selected GO terms. The Biological Process (BP) category is shown in cyan, the Molecular Function (MF) category is shown in yellow, and the Cellular Component (CC) category is shown in magenta. See Additional File 5 for complete GO term analysis of downregulated genes.

We used motif finding software to scan the 250bp regions upstream of the transcription start sites of the downregulated genes, and identified a putative transcription factor binding site (Motif D1, Figure 5A). This motif is homologous to the identified Hr3 binding site (p = 0.005; Figure 5A). However, we find that Motif D1 is not enriched among downregulated genes in comparison with our unchanged control gene set (odds ratio 1.14, p = 0.826), and that the presence of Motif D1 does not have a significant effect on the rank-product among downregulated genes (W = 97, p = 0.811; Figure 5B), suggesting that this site is likely not mediating the downregulation observed following infection.

**Figure 5.**
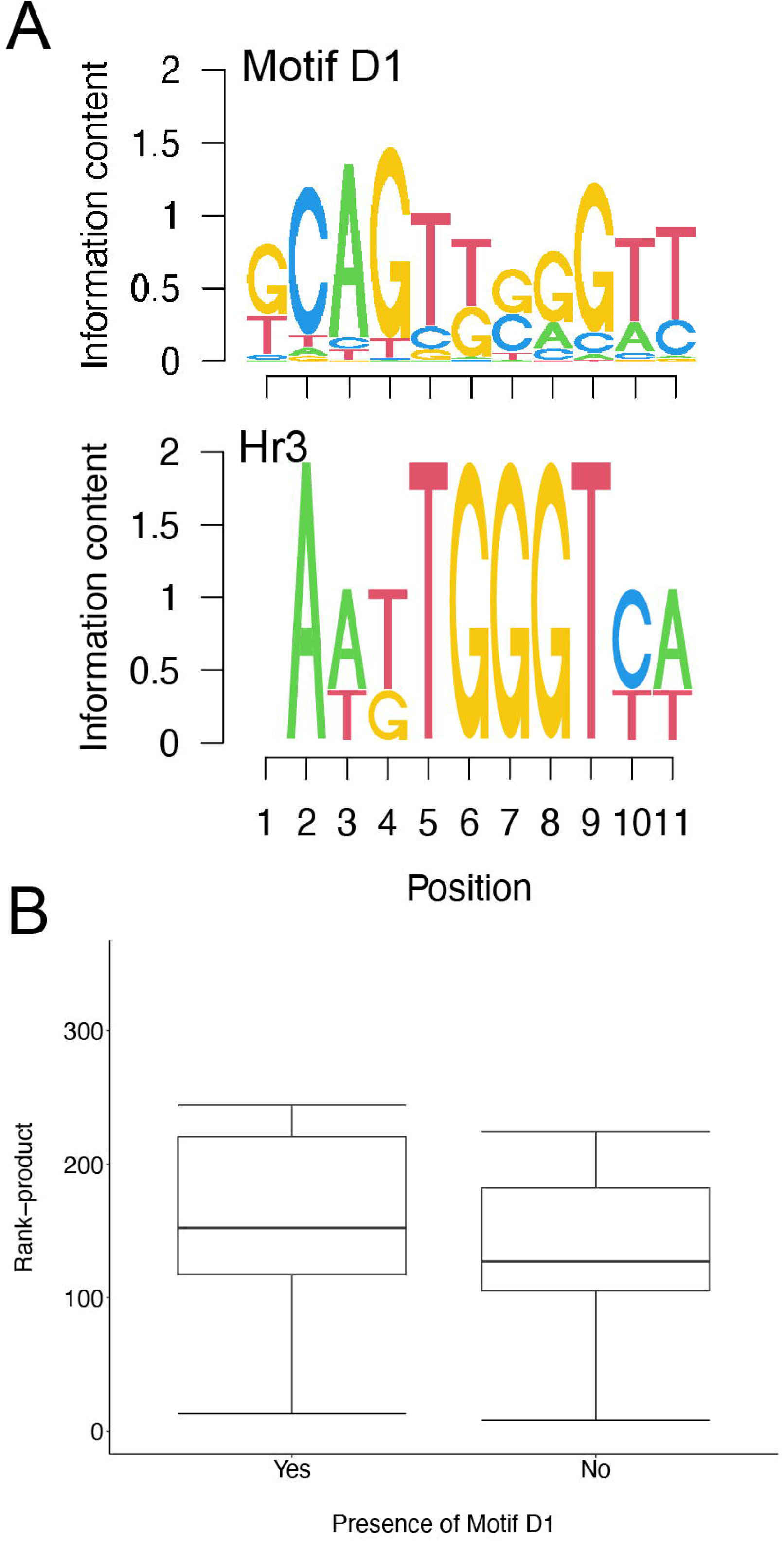
Motif associated with downregulated genes. *De novo* motif finding identified 1 motif (Motif D1) that is enriched in the upstream sequences of the induced genes. The consensus motif is represented as a sequence logo (A, top). Motif matching identifies Motif D1 as being significantly similar to the Hr3 binding site (A). (B) Box-whisker plot showing the distribution of rank-products for the downregulated genes with and without Motif D1. The presence of Motif D1 does not impact the rank-product distribution.

## Discussion

Our findings have uncovered a subset of *Drosophila melanogaster* genes whose expression is altered following infection by a range of pathogens. These genes are likely mediating known responses to infection, for instance we find that known immune response genes predominate among the induced genes on our list. Many of the downregulated genes we identified are linked to metabolism, feeding, development, and reproduction; processes which are all altered following infection. Additionally, our analysis has uncovered a putative transcriptional mechanism that regulates gene expression following infection with diverse pathogens.

In the absence of *in vivo* experimental data, we are unable to draw conclusions about the roles of the genes we’ve identified, however our bioinformatic and meta-analyses do allow us to generate interesting hypotheses for future testing. We believe that our lists of induced and downregulated genes can provide insight in host immunity, particularly due to the presence of genes with experimentally defined functions that align with observed immune response mechanisms as highlighted below; however these roles need to be empirically tested in future studies. Hopefully our analyses have provided an interesting list of candidate genes whose study can begin to unravel important immune mechanisms.

Intriguingly, we find that many specific immune effector encoding genes are induced following infection by diverse pathogens. For instance, antimicrobial and AMP-like peptides are not known to play a role in the antiparasite immune response, and yet are induced following parasitoid wasp infection (Table 4) [19, 88, 89]. A possible model to explain this observation is that the *Drosophila* genome encodes a conserved set of immune effector genes that act against all types of pathogen infection. This model is unlikely given the long history of findings suggesting that specific immune effectors are used to target distinct pathogens [24, 90]. Additionally, multiple studies have demonstrated that selection for flies with resistance to a particular pathogen does not translate to cross-resistance to additional pathogens [91, 92]. Specifically, flies selected for resistance to the parasitoid *Asobara tabida* do not show increased resistance to bacterial or fungal pathogens [91], and fly lines selected for resistance to the bacterial pathogen *Bacillus cereus* do not display cross-resistance to Sigma virus [92].

Instead, our findings may suggest a second model in which sensing pathogen infection leads to the induction of genes that play a role in surveillance and resistance to possible coinfecting pathogens. Coinfections are commonly observed in natural populations of various species [93–95]. It has been demonstrated that coinfection can lead to decreased host resistance in nature and laboratory experiments [95–97], and negatively impact host health [94, 98]. These previous findings suggest that avoidance of coinfection would increase host fitness, and support a model in which infection by any pathogen may provoke a generalized prophylactic response against coinfection alongside the specific response to the primary pathogen. Indeed, we have identified a large number of induced genes that encode immune recognition proteins and regulators of the Toll, IMD, and JAK-STAT immune signaling pathways, along with immune effectors that target distinct classes of pathogens. These pathways are among the main immune response pathways in *Drosophila*, and in combination with the breadth of induced immune effectors, our findings suggest that infected flies are primed to respond to the possibility of coinfection.

This model is also supported by previous findings. For instance, AMP expression is seen at early time points following parasitoid infection [19, 88], but little to no expression of antimicrobial immune effectors is observed at late time points following parasitoid infection [89]. These findings may make sense in the light of the coinfection prevention model. In nature, *Drosophila* larvae are found in the microbe-rich environment of decaying fruit [99]. Parasitoid infection of *Drosophila* larvae results in the wasp ovipositor puncturing a hole in the larval cuticle; this wound will be healed, however the healing of epidermal wounds can take several hours [100, 101]. Immediately following parasitoid infection, and before healing is complete, the wound can therefore provide a readily available infection route for environmental microorganisms. The expression of antimicrobial factors and surveillance for any surviving microbes may therefore play an important role in pre-empting this possible route of coinfection.

Additional support for this model is provided by an in-depth time course study of the transcriptional response to IMD pathway activation [102]. In this study, stimulation of the IMD pathway resulted in the expression of Toll pathway regulated genes, including *Bom* family genes, and stress response genes, including *TotM,* all of which we also identified in our meta-analysis. Interestingly, the high resolution time course provided by this study illustrates that the Toll and stress response genes were induced as part of an early and transient response to IMD pathway stimulation, in contrast to the sustained transcription of known IMD responsive genes [102]. This pattern would fit with our expectations under a coinfection prevention model, in which immune stimulation simultaneously triggers a specific response against the identified pathogen (IMD pathway genes) and leads to the production of a temporary prophylactic state (Toll and stress response genes) to guard against possible coinfection.

Using *de novo* motif finding analysis, we identified putative binding sites for NFκB and GATA family transcription factors. The degree of gene induction in our meta-analysis correlates with the presence of these factors, and suggests a possible synergistic relationship between them. NFκB and GATA factor activity have been linked to immune responses, and have been previously demonstrated to work in concert to promote immune gene expression [103], supporting our idea that these factors may be underlying the response to infection by diverse pathogens. The *Drosophila* genome encodes 3 NFκB family genes (*Dif, dl, Rel*), all of which have been previously linked to immunity, and 5 GATA factors (*pnr, srp, grn, dGATAd, dGATAe*) of which *srp* and *dGATAe* have been previously linked to immune responses [68–71, 104]. The difficulty in distinguishing between paralog-specific binding site motifs within these families leaves us unable to speculate whether the response is driven by a particular family member, or whether multiple members may play a role. Cross-regulation of gene expression has been observed between the NFκB-dependent Toll (*Dif* and *dl* dependent) and IMD (*Rel* dependent) pathways [105], perhaps suggesting some redundancy between NFκB factors. In order to build a model of transcriptional regulation the role of individual factors must still be tested experimentally, and the results may help in understanding the immune response to infection.

While downregulated genes are often overlooked in studies of gene expression, they may still provide insight into the process being studied. Our meta-analysis identified a small number of transcripts that are downregulated following infection by diverse pathogens. The functions of these genes suggest that they may be playing a role in the switch to an altered metabolic state following infection. Infected flies display altered feeding behaviour, and prioritize using energy resources for immunity ahead of development or reproduction [75, 76, 82, 106]. Accordingly, we find that genes previously linked with these processes are downregulated in infected flies. The further study of these genes may shed light on the largely unknown mechanisms underlying the life history tradeoffs induced by pathogen infection.

## Conclusions

Our meta-analysis has identified 93 genes whose transcript levels are significantly altered following infection by diverse pathogens. Analysis of the experimentally determined and predicted functions of the proteins encoded by these genes suggests that they may play a role in immune function, immune metabolism and infection induced life history tradeoffs. Follow up studies on the roles of these genes following infection will be necessary to verify their importance and will likely improve our understanding of conserved immune functions.

## Methods

### Drosophila melanogaster genome data

Gene expression data were imported from the National Center for Biotechnology Information (NCBI) Gene Expression Omnibus (GEO) database (https://www.ncbi.nlm.nih.gov/geo/) [107] and the Berkeley *Drosophila* Genome Project (BDGP) (https://www.fruitfly.org/expression/immunity/index.shtml). Accession numbers and other metadata are listed in Table 1.

Gene expression data were then pre-processed before meta-analysis. First, gene identifiers were converted to the most recent FlyBase gene identification number (FBgn) using the FlyBase Upload and Validate IDs tool (version FB2021_01) [108]. Second, the gene expression datasets were filtered to remove any genes that are not represented in all of the datasets. This step resulted in the identification of 10,818 common genes that were retained for subsequent analysis. Finally, gene expression fold change values were log_2_ transformed wherever necessary to be used as input for meta-analysis (next section). Data for the *D. melanogaster* genome (release 6.38) and individual gene reports were accessed through FlyBase (version FB2021_01) [108].

### Meta-analysis of gene expression studies

Meta-analysis of immune gene expression studies was performed in the R statistical computing environment [109], using the RankProd package [43, 44]. The log_2_ fold change for each gene in each dataset was used as the input, and the rank and rank-product (RP) were calculated for each gene. Significance was determined using the estimated percentage of false prediction (pfp) with a threshold of 0.05. Genes with significantly altered expression are listed in Additional Files 1 (62 upregulated genes) and 5 (31 downregulated genes). A control set of 62 genes with unchanged expression was selected from the RP result as listed in Additional File 4.

### Chromosomal distribution

The chromosomal location of each gene identified as significantly differentially expressed in the meta-analysis was retrieved from FlyBase (version FB2021_01). The proportion of identified genes found on each chromosomal arm was compared to the proportion of all genes for that arm using a 2-sample test for equality of proportions with continuity correction in R.

### Gene locus uniformity and clustering

To determine the uniformity of gene spacing across *D. melanogaster* chromosome arms, the up- and down-regulated gene lists were used as input for the Cluster Locator webserver (http://clusterlocator.bnd.edu.uy/ accessed 4/24/2021) [110] using default parameters. Uniformity is tested using a two-sided Kolmogorov-Smirnov test. Clusters were identified using the Cluster Locator webserver with the default max-gap value of 5. Gene clustering is statistically tested by comparing clustering of the input lists with randomly selected lists.

### Motif finding

Putative transcription factor binding motifs were predicted using iMotifs [62, 111]. The 250bp of sequence immediately upstream of each gene of interest was downloaded from FlyBase, and these sequences were used as input to iMotifs. These predicted motifs were then mapped onto the input and unchanged control sequences using the FIMO tool [112]. To identify the likely transcription factor interacting with the discovered motifs, the motifs were compared with experimentally validated *D. melanogaster* transcription factor binding sites using the TomTom Motif Comparison Tool [63, 113] with a significance threshold of p < 0.05. Position-weight-matrices for the identified transcription factor binding sites were accessed through the OnTheFly database [114]. OnTheFly accession numbers: GATAe: OTF0433.1; dl: OTF0107.2; Hr46/Hr3: OTF0227.1.

### Statistics

All statistical tests were performed in R using the base stats package and graphs were produced using ggplot2 [115].

## Additional Files

**Additional File 1**: All significantly induced genes from the RankProduct analysis listed by FBgn and gene name. The following statistics are included in the table: RP is the calculated rank-product; FC is the fold change and logFC is the log_2_ fold change between infected and naïve across the datasets; pfp is the estimated percentage of false prediction; P.value is the uncorrected p value. Significance was determined using a pfp cutoff of 0.05.

**Additional File 2**: All significantly downregulated genes from the RankProduct analysis listed by FBgn and gene name. The following statistics are included in the table: RP is the calculated rank-product; FC is the fold change and logFC is the log_2_ fold change between infected and naïve across the datasets; pfp is the estimated percentage of false prediction; P.value is the uncorrected p value. Significance was determined using a pfp cutoff of 0.05.

**Additional File 3**: All significantly enriched Gene Ontology terms among genes identified as induced by the RankProduct analysis

**Additional File 4**: Genes identified as unchanged from the RankProduct analysis and used as a representative background set for motif analysis. Genes are listed by FBgn and gene name. The following statistics are included in the table: RP is the calculated rank-product; FC is the fold change and logFC is the log_2_ fold change between infected and naïve across the datasets; pfp is the estimated percentage of false prediction; P.value is the uncorrected p value.

**Additional File 5**: All significantly enriched Gene Ontology terms among genes identified as downregulated by the RankProduct analysis

## Declarations

### Ethics approval and consent to participate

Not applicable

### Consent for publication

Not applicable

### Availability of data

The datasets analysed in the current study are available through the Gene Expression Omnibus (https://www.ncbi.nlm.nih.gov/geo/ study accessions: GSE37708, GSE2828, GSE42726, GSE25522, GSE8938, GSE5489, GSE31542), and through the fruitfly.org Expression Database (http://www.fruitfly.org/expression/immunity/).

### Authors’ contributions

Conceptualization: NTM; Investigation: ALW, JH, BA, NB, NL, PKR, DF, NTM; Analysis: ALW, JH, BMA, NB, NL, PKR, DF, NTM; Supervision: NTM; Funding Acquisition: NTM; Writing – Original Draft: NTM; Writing – Review & Editing: ALW, JH, BMA, NB, EL, PKR, DF.

## Acknowledgements

We would like to acknowledge Dr. Alysia Vrailas-Mortimer and members of the Mortimer Cellular Immunology Lab for discussion of results.

